# Dynamic clamp constructed phase diagram of the Hodgkin-Huxley action potential model

**DOI:** 10.1101/766154

**Authors:** Hillel Ori, Hananel Hazan, Eve Marder, Shimon Marom

**Affiliations:** Technion – Israel Institute of Technology, Haifa 32000, ISRAEL; Brandeis University, Waltham, MA 02454-9110

## Abstract

Excitability – a threshold governed transient in transmembrane voltage – is a fundamental physiological process that controls the function of the heart, endocrine, muscles and neuronal tissues. The 1950’s Hodgkin and Huxley explicit formulation provides a mathematical framework for understanding excitability, as the consequence of the properties of voltage-gated sodium and potassium channels. The Hodgkin-Huxley model is more sensitive to parametric variations of protein densities and kinetics than biological systems whose excitability is apparently more robust. It is generally assumed that the model’s sensitivity reflects missing functional relations between its parameters or other components present in biological systems. Here we experimentally construct excitable membranes using the dynamic clamp and voltage-gated potassium ionic channels (Kv1.3) expressed in *Xenopus* oocytes. We take advantage of a theoretically derived phase diagram, where the phenomenon of excitability is reduced to two dimensions defined as combinations of the Hodgkin-Huxley model parameters. This biological-computational hybrid enabled us to explore functional relations in the parameter space, experimentally validate the phase diagram of the Hodgkin-Huxley model, and demonstrate activity-dependence and hysteretic dynamics due to the impacts of slow inactivation kinetics. The experimental results presented here provide new insights into the gap between technology-guided high-dimensional descriptions, and a lower, physiological dimensionality, within which biological function is embedded.

---

The canonical Hodgkin-Huxley model of excitability [1] consists of four dynamical variables (membrane voltage and three protein state variables) and more than ten parameters. Several of the parameters represent actual, measurable physical entities (membrane capacitance, ionic concentrations inside and outside the cell, densities of ionic channel membrane proteins). Other parameters shape the six exponential functions relating membrane voltage to probabilities of transitions between protein states [2–4]. The considerable sensitivity of the model to parametric variations – especially protein densities and kinetics – is not on par with the robustness of many biological systems, as revealed in experiments showing a high degree of resilience to variation in values of measurable parameters [5–8]. Since the Hodgkin-Huxley model is biophysically solid, it is generally assumed that its sensitivity reflects functional relationship between parameters [7]. Hence, a low dimensional representation of Hodgkin-Huxley parameter space within which the seemingly complicated and parameter-sensitive system becomes tractable, would help understand how a biological system could control its state and adapt to changes using a simple and physiologically relevant process.

Recently, a biophysically oriented parametrization of the Hodgkin-Huxley model was introduced [4], offering a framework for understanding control of excitability amid changes in protein densities and their kinetics. In the resulting phase diagram, the “excitability status” of a given Hodgkin-Huxley realization is determined by rational functions fully defined in terms of Hodgkin-Huxley parameters along two physiological dimensions: structural and kinetic. The structural dimension (denoted S) is a measure for the relative contribution of maximal exciting (i.e. sodium) conductance. The kinetic dimension (denoted K) is a measure of the relative contribution of restoring voltage dependent rate functions that pull the membrane back to its hyperpolarized potential (i.e., closure of sodium channels, opening of potassium channels). Thus, a point in the S–K phase diagram represents many different possible sets of the model parameters, explicit instantiations of a Hodgkin-Huxley model that give rise to similar functional outcome. Examined in the S–K plane, the Hodgkin-Huxley model reveals order that is impossible to detect at higher dimensional representations. The three different excitability statuses are clustered in “phases”: non-excitable, excitable, and oscillating (depicted in the left side panel of Figure 1). Note that the S–K phase diagram is different from a phase space where each point depicts a unique membrane state of a given instantiation, and where lines connecting such states depict phase portraits, a trajectory in time [e.g., 9–12].

**Figure 1:**
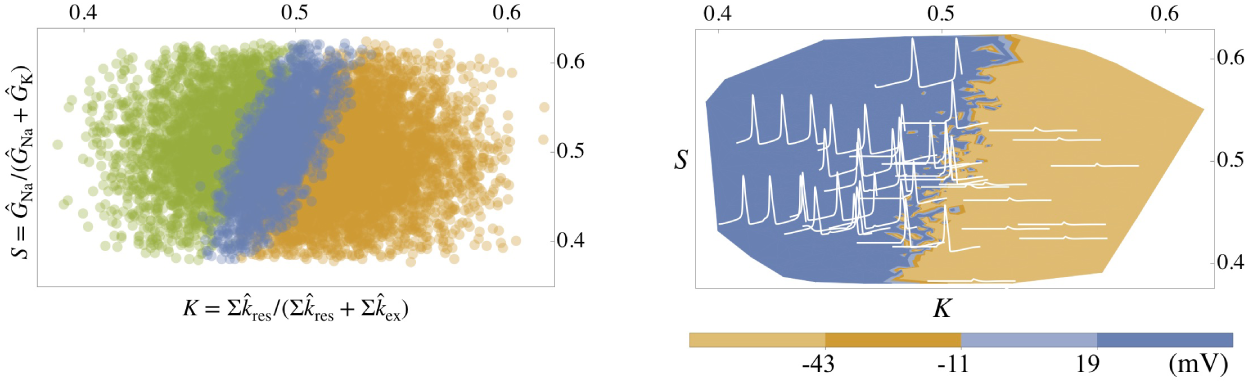
Structural–Kinetic (S–K) phase diagram of excitability status in the Hodgkin-Huxley model. **Left:** 10000 realizations of a full Hodgkin-Huxley model, following the procedure described in Ori *et al*. (2018). *Ĝ*_X_ is the maximal membrane conductance to ion X relative to the Hodgkin-Huxley standard model values 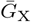, where 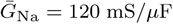 and 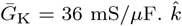 is a linear scalar of ionic channel transition rate function. The sub-scripts of 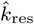 and 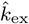 epict transition rate functions contributing to restoring and exciting forces: *α*_n_(*V*) and *β*_m_(*V*) for the former, and *α*_m_(*V*) and *β*_n_(*V*) for the latter. Parameters (maximal sodium and potassium conductance, leak conductance, membrane capacitance and the six rate equation functions) vary randomly and independently over a ±0.25 range compared to their values in the original Hodgkin-Huxley model. The resulting membrane responses to above-threshold stimuli are classified (different colors) to three excitability statuses: excitable (2225, blue), not excitable (4884, orange), and oscillatory (2891, green). **Right:** data set of the left panel replotted with colors depicting response peak amplitude clusters, classified to four bins indicated in the horizontal color bar.

The present study aims to experimentally uncover the S–K phase diagram, with actual biological components rather than mathematical modeling. The challenge is this: By definition, a cell at a specific moment in time is an instantiation of one point in the S–K phase diagram, one set of parameters. It is therefore impossible to systematically cover the S–K plane of a given biological cell (neuron, cardiac myocyte, etc.) by manipulating the kinetic and structural features of its constituents, the ionic channel proteins. Collecting data from many individual cells of similar type would not help, because similar types have a tendency to be residents of same phase (e.g., cardiac myocytes are mostly oscillating, and cortical neurons are mostly excitable but not oscillatory). Here we face the challenge of systematic sampling the Hodgkin-Huxley S–K phase diagram by combining the established methodology of heterologous expression of channel proteins in *Xenopus* oocytes [13, 14] and hard real-time dynamic clamp [15–18]. With this approach we experimentally reconstruct the first phase transition in the S–K diagram: the transition between the non-excitable and excitable phases (depicted in the right-side panel of Figure 1). Moreover, we show that directional “walk” within the S–K plane exposes hysteresis in the organization of the phase diagram, which we explain in terms of channel protein slow and activity-dependent gating that potentially enables control of excitability amid parametric variation [3, 4].

## Results & Discussion

The *Xenopus* oocyte protein expression system is frequently used in physiological studies of ionic channels; several fundamental studies of ion channel structure-function relations were made using this simple and experimentally elegant system [e.g., 19–21]. For all practical purposes the oocyte is an ideal electrophysiological ghost: it is a large and spherical (i.e., iso-potential) leaky capacitor, it does not express significant voltage-sensitive membrane conductances, and it readily expresses functional conductances following injection of ionic channel coding mRNAs. Wedding heterologous *Xenopus* oocyte expression with dynamic clamp makes it possible to split the system’s components between those that are biologically expressed in the oocyte membrane and those that are computationally expressed in the dynamic clamp algorithm. Figure 2A demonstrates the efficacy of this approach in generating a bio-synthetic excitable system: An oocyte is impaled by two sharp electrodes. Signals from the voltage measuring electrode are read by a real-time processor that calculates Hodgkin-Huxley sodium and potassium currents, feeding the sum of these currents back to the oocyte through the current injecting electrode. The oocyte contributes membrane capacitance and leak conductance, the linear components; the dynamic clamp algorithm contributes sodium and potassium voltage-dependent conductance, the non-linear components. The nature of the system’s response (bottom of 2A) depends on the Hodgkin-Huxley parameters implemented in the dynamic clamp algorithm.

**Figure 2:**
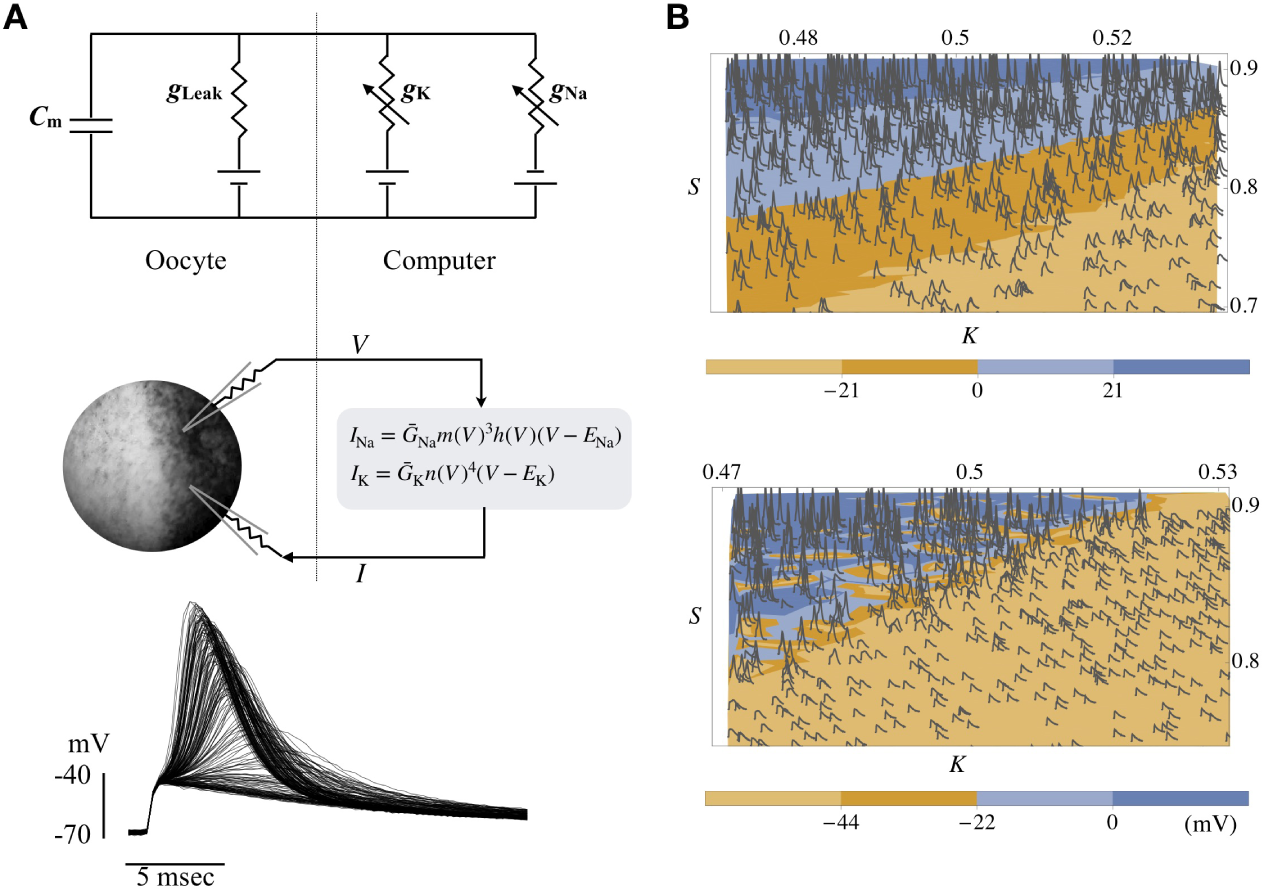
Dynamic clamp constructed phase diagram of excitability in naive (i.e., no channel mRNA injected) *Xenopus* oocyte. **A:** Equivalent electrical circuit of the basic system configuration (top and middle panels). An oocyte is impaled by two sharp electrodes. Signals from the voltage measuring electrode are read by a real-time processor that calculates Hodgkin-Huxley sodium and potassium currents, feeding the sum of these currents back to the oocyte through the current injecting electrode. The oocyte contributes membrane capacitance and leak conductance; the dynamic clamp algorithm contributes sodium and potassium voltage-dependent conductance. Excitability may be induced (bottom) upon activation of the dynamic clamp algorithm. The nature of the system’s response to stimuli (in this case 0.5 msec, 10*µ*A; see Methods) depends on the Hodgkin-Huxley parameters implemented in the dynamic clamp algorithm (bottom panel: *Ĝ*_Na_ =[0,10], 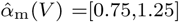). **B:** S–K phase diagrams. The ranges of S and K accessible for scanning by dynamic clamp differ between experiments; they are dictated, mainly, by leak and capacitance contributed by the oocyte and affected by the quality of electrodes-membrane interactions. In both experiments, soft but relatively well-defined borders separate non-excitable from excitable phases (colors depict peak response amplitude clusters, classified to four bins indicated in horizontal color bars). The diagrams were constructed under different conditions: in the top panel the parameters of sodium and potassium conductance were taken from the Hodgkin-Huxley canonical model. In the bottom panel, the potassium conductance parameters are those of the Kv1.3 channel [22]. Both experiments were conducted in the same oocyte.

Following Ori *et al*. (2018), the structural dimension (S) is defined as *Ĝ*_Na_*/*(*Ĝ*_Na_+ *Ĝ*_K_) where *Ĝ*_X_ is the maximal membrane conductance to ion X relative to the Hodgkin-Huxley standard model values 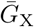, where 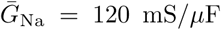 and 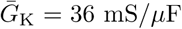. For instance, *Ĝ*_Na_ = 1.25 stands for 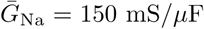. The kinetic dimension (K) is defined as 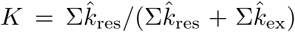, where 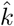 is a linear scalar of ionic channel transition rate function. For instance, the expression 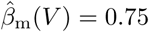 stands for 0.75*β*_m_(*V*). The subscripts of 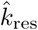 and 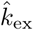 depict transition rate functions contributing to restoring and exciting forces: *α*_n_(*V*) and *β*_m_(*V*) for the former, and *α*_m_(*V*) and *β*_n_(*V*) for the latter.

To experimentally construct an S–K phase diagram in the dynamically-clamped *Xenopus* oocyte, a random list of 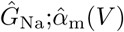 pairs was generated ([0,10];[0,2], respectively). These, in turn, were translated to S–K values, as *S* = *Ĝ*_Na_*/*(*Ĝ*_Na_ + 1) and 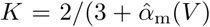. The excitability status of the oocyte for each instantiation of an S–K value was determined by its response to a single, constant amplitude current stimulus (0.5 msec; see Methods). The resulting phase diagrams of two experiments are presented in Figure 2B. Similar to the theoretical phase diagram of Figure 1 (right side panel), the plot of Figure 2B shows well-structured S–K planes with soft but relatively well-defined borders that separate non-excitable from excitable phases (colors depict response amplitude clusters, classified to four bins indicated in horizontal color bars). In the top panel of Figure 2B, the parameters of sodium and potassium conductance were taken from the Hodgkin-Huxley canonical model. In the bottom panel, the potassium conductance parameters are those of the Kv1.3 channel [22–24].

To validate the reduction to an S–K space, we take the above experimental system a step further by “relocating” the voltage-dependent potassium conductance from the dynamic-clamp algorithm into the biological domain (Figure 3A). This is achieved by injection of mRNA that codes the voltage-dependent Kv1.3 potassium channel. Within a few days, the channels are extensively expressed in the oocyte membrane. As demonstrated in Figure 3A (lower panel), upon activation of the dynamic clamp, excitability emerges with biological capacitance, leak and potassium conductance, while the sodium conductance and its related kinetics are expressed computationally. [Our attempts to implement the inverse experimental condition, where sodium conductance is biologically expressed, did not succeed; the expression of sodium conductance was too weak to support full-blown excitability.]

**Figure 3:**
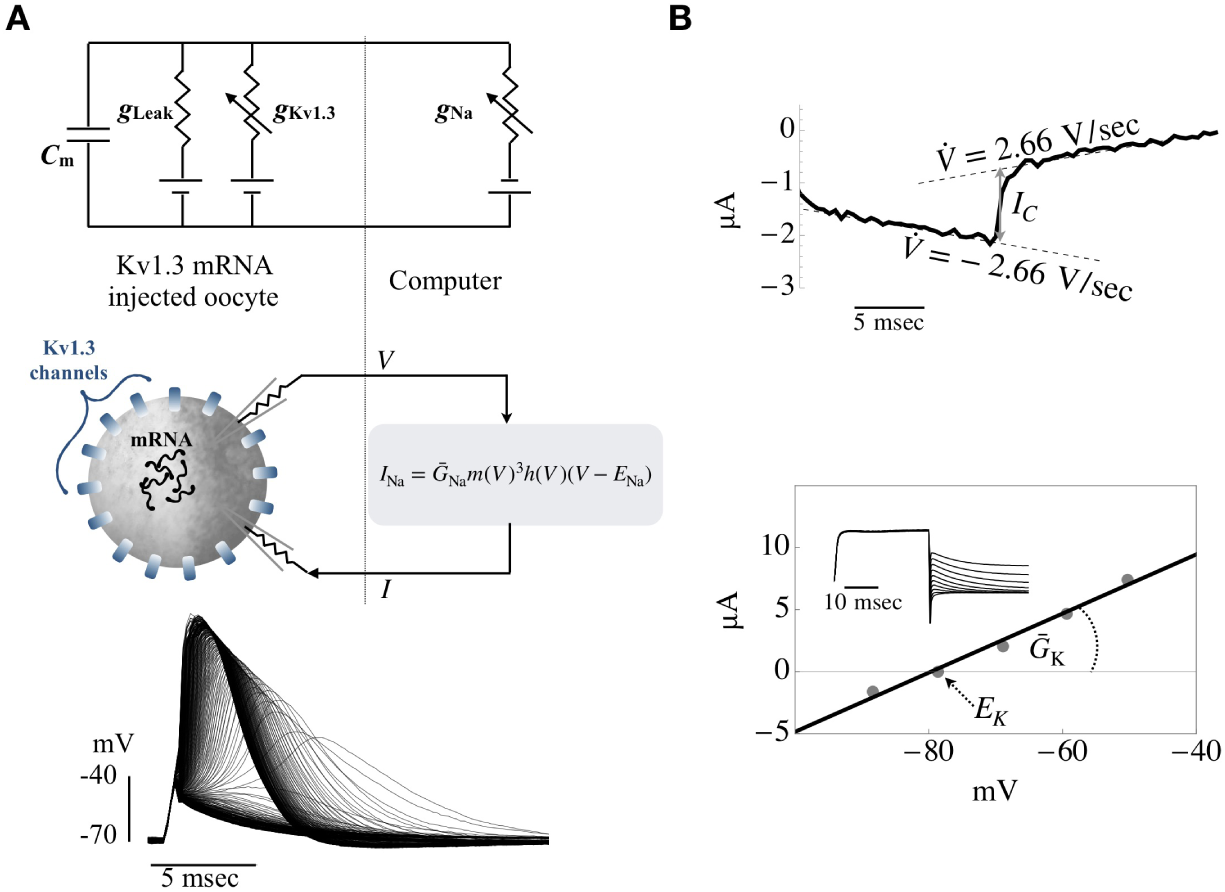
Dynamic clamp and excitability in Kv1.3 mRNA injected *Xenopus* oocyte. **A:** Taking the experimental system described in Figure 2 a step further by “re-locating” the voltage-dependent potassium conductance from the dynamic-clamp algorithm into the biological domain (top panel). This is achieved by injection of mRNA that codes the voltage-dependent Kv1.3 potassium channel. Within a couple of days, the channels are extensively expressed in the oocyte membrane. Upon activation of the dynamic clamp, excitability emerges with biological capacitance, leak and potassium conductance, while the sodium conductance and its related kinetics are computationally expressed. The nature of the system’s response depends on the Hodgkin-Huxley parameters implemented in the dynamic clamp algorithm (bottom panel: *Ĝ*_Na_ =[0,16], 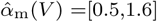). **B:** Maximal potassium conductance, potassium Nernst potential and membrane capacitance are estimated by standard voltage-clamp procedures. Top: Cell capacitance is estimated from the capacitive current step upon instantaneous switch from −2.66 V/sec to +2.66 V/sec (in a range between −70 to −110 mV). Bottom: Tail current protocol (inset) to establish potassium Nernst potential based on current reversal; depolarizations to +40 mV followed by hyperpolarization to different values. 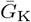 was estimated from maximal current and Nernst potential, and validated using tail current data.

Since there is no “standard” model for excitability with Kv1.3 conductance, we express the structural dimension in terms of actual maximal conductance normalized to membrane capacitance; thus, 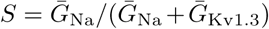. Maximal potassium conductance 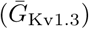, potassium Nernst potential and membrane capacitance are estimated by standard voltage-clamp procedures (Figure 3B; and see Methods section). We take the voltage-dependent rate functions of the Kv1.3 conductance [22, 24] as reference to express the kinetic dimension, hence 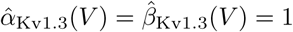.

The phase diagrams of three different experiments are presented in Figure 4, where S–K planes with well-defined borders that separate non-excitable from excitable phases validate the reduction of the original Hodgkin-Huxley high-dimensional parameter space to the lower S–K phase diagram. Borders that separate non-excitable from excitable phases in 12 different experiments are summarized in the bottom-right panel.

**Figure 4:**
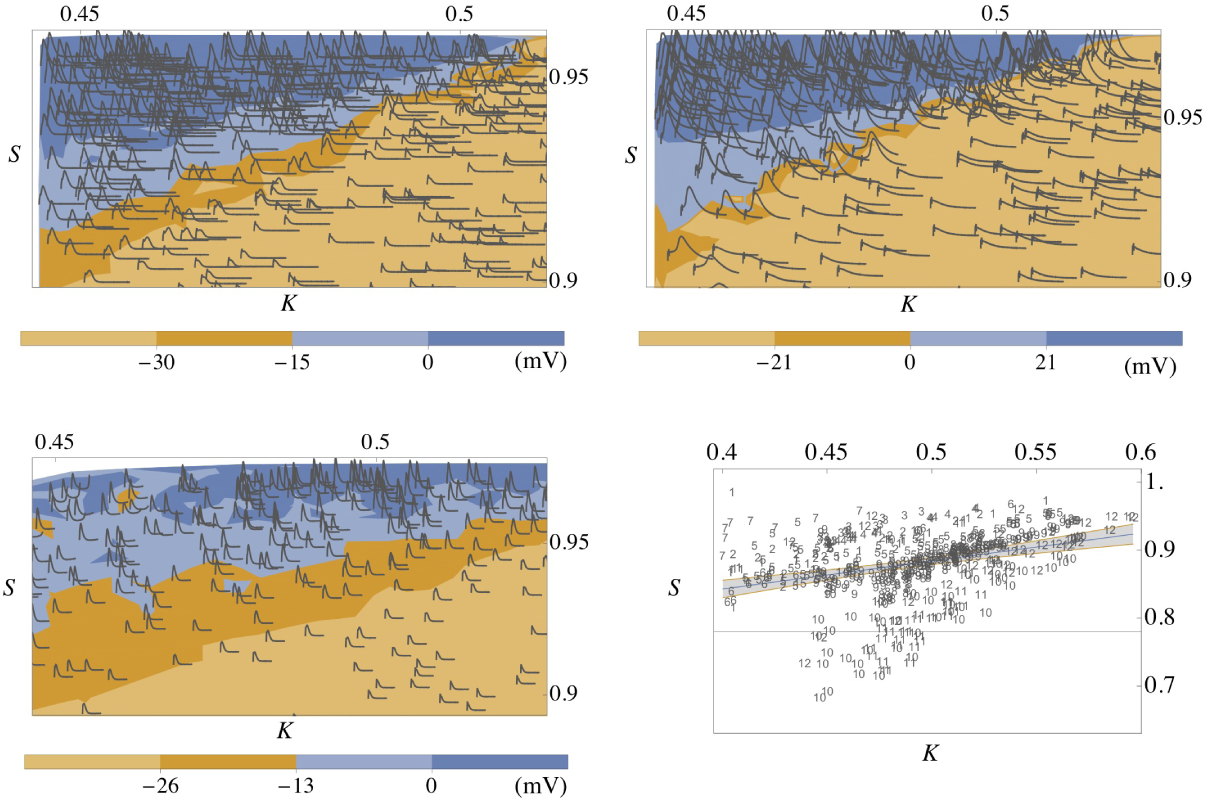
Dynamic clamp constructed phase diagram of excitability in Kv1.3 injected oocytes. Top two panels and bottom-left panel demonstrate phase diagrams of three different experiments, where biological capacitance, leak and potassium conductance are contributed by the oocyte, while the sodium conductance and its related kinetics are “computationally expressed” in the dynamic clamp algorithm. Note S–K planes with relatively well-defined borders that separate non-excitable from excitable phases. Bottom-right: Summary of 12 experiments, showing that a positive slope diagonal separating non-excitable from excitable phases (demonstrated in the three other panels) is consistent across experiments. To this end, a histogram of response amplitudes was generated for each experiment. Of the 9889 responses in all 12 experiments, 529 responses were identified within an intermediate range [0.2, 0.8] of response amplitudes (the amplitude of the passive response to the stimulus was defined as zero). The S–K coordinates of these 529 responses are plotted in the bottom-right panel, together with a fitted straight line (*S* = 0.7 + 0.4*K*; 99% confidence bands for mean predictions). Numerical symbols depict the different experiments. The slopes of the individual lines fitted separately for each of the 12 experiments are: 0.66, 0.62, 0.62, 0.75, 0.58, 1.01, 0.49, 1.46, 0.67, 1.25, 2.2, 0.73. Note that these slopes are significantly less steep compared to the slope of the Hodgkin-Huxley model reported by Ori *et al*. [4].

The power of the parametrization may be further appreciated by observing multiple instantiations of the same S–K coordinate, demonstrating that the outcome is quite resilient to the actual set of parameters used to determine a given coordinate (Figure 5). Even delicate response features (e.g., the post-stimulus sub-threshold depolarization in the right-most traces of the bottom panel), not accounted for in the construction of the theory [4], are nicely caught by the S–K coordinates.

**Figure 5:**
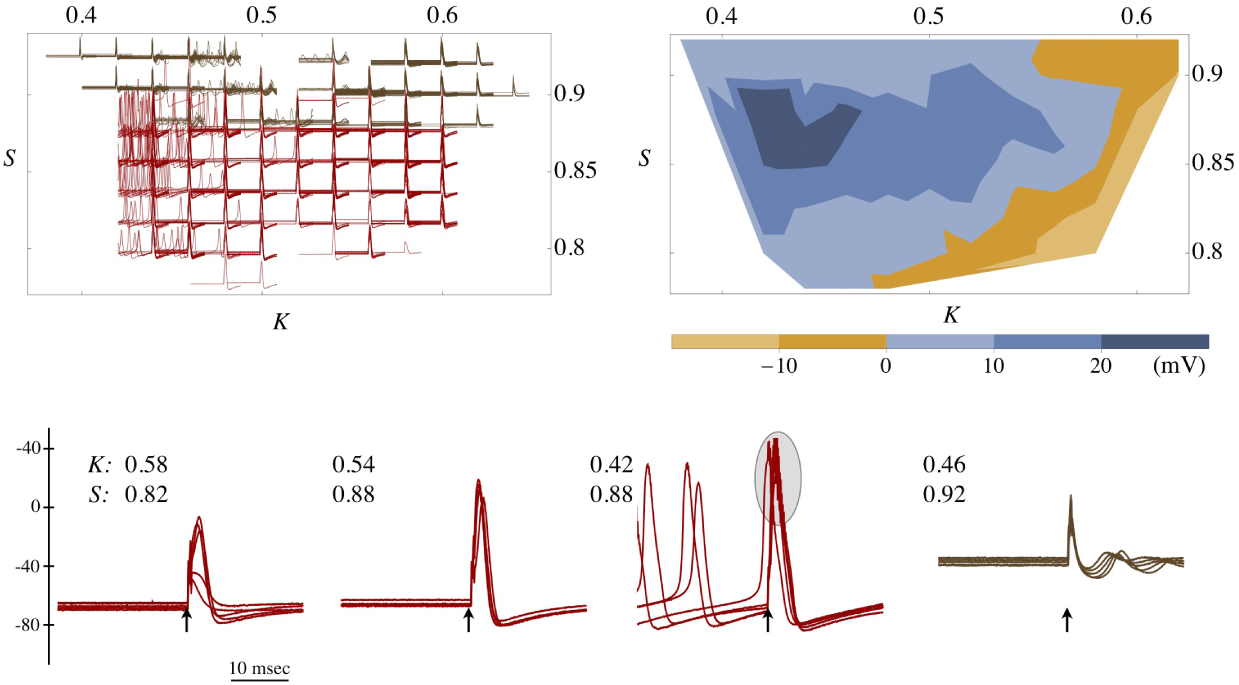
The adequacy of the S–K parametrization estimated by observing multiple instantiations of the same S–K coordinate. A Kv1.3 mRNA injected oocyte 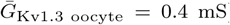 is coupled to a dynamic clamp algorithm where the Hodgkin-Huxley sodium, potassium and leak conductance (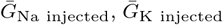 *G*_Leak injected_, respectively) and their related kinetics are computationally expressed. A set of 3000 instantiations was prepared, within the following ranges: 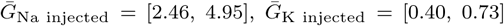 OR set to zero (see below), 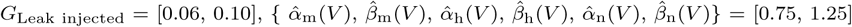. The membrane response to a 0.5 msec stimulus (14.5 *µ*A) was recorded. Different instantiations were delivered at a rate of 5/sec. Assuming 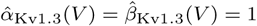 and considering both potassium conductance (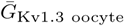 and 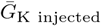), the values of S and K for each instantiation was calculated and binned to 0.02 resolution. For clarity, the total of 3000 traces was down-sampled by a factor of 3, and each of the sampled responses is plotted in its corresponding S–K coordinate bin in the **top left** panel. The brown colored traces are those corresponding to cases where 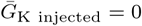; the red colored traces are those corresponding to cases where 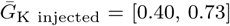. **Top right**: contour plot of mean peak amplitudes for all binned S–K coordinates (3000 spikes). **Bottom** four panels: Examples of several responses in 4 of the S–K coordinates (arrows depict stimulus time; traces were smoothed by moving average of 120 *µ*sec). The responses are fairly resilient to the actual set of parameters used to determine a given coordinate. Even delicate response features (e.g., the post-stimulus sub-threshold depolarization in the right-most traces), not accounted for in the construction of the theory, are nicely caught by the S–K coordinates. Note: “ringing” in several of the fast-high amplitude spikes (depicted by gray oval in one of the bottom panels) are due to limits imposed by the rate of the real time processor and/or the current injection device (see methods).

Over the years since Hodgkin and Huxley presented their canonic model, physiologists have continued to explore the complex behavior of voltage-gated ion channels. In particular, not all transition rates are voltage-dependent, and their characteristic time scales span a wide range that extends from sub-milliseconds to minutes [25–29]. The results presented in Figures 2, 4 and 5 are limited to the short, millisecond timescale, hence slow activity-dependent effects could not be detected. But slow channel protein gating and its impacts on response dynamics might be exposed by “traveling” in a directional manner within the S–K diagram; for instance, by moving up-and-down along a ramp within the diagram. The kinetics of Kv1.3 are particularly relevant in this context, as they involve voltage and state-dependent transition rates, and a mix of slow and fast reactions spanning a wide range of time scales [22, 24]. Moreover, these kinetics of Kv1.3 were suggested to have significant impacts on excitability on longer time scales [30–33]. The left panel of Figure 6 shows membrane responses of a Kv1.3 expressing oocyte as a function of S–K coordinates in a directed walk within the diagram. A gradual (up and down ramp; total trajectory 450 sec) change is implemented in 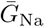 and 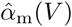, dynamic clamp parameters of Hodgkin-Huxley sodium conductance. The spikes are plotted over an S–K diagram constructed for that same oocyte. Each trace depicts a membrane response to a short above-threshold current stimulation. As the system is moved toward the excitable phase (top-left corner of the diagram), the oocyte responds to the stimulus with a self-propelled depolarization that becomes a fully blown action potential. At some point, as might be expected, the structural exciting force (S) is so high and the kinetic restoring force (K) is so low (0.95, 0.42, depicted by arrow in the right panel) that the membrane cannot hyperpolarize back to resting potential, and remains stuck in a depolarized, not excitable value. Upon return, clear hysteresis is revealed, reflecting recovery of the Kv1.3 from long lasting inactivation. The insets to the right panel of Figure 6 show two repetitions of the same ramp protocol in another oocyte, demonstrating reversibility of the hysteresis phenomenon. Such hysteresis is not seen with the Hodgkin-Huxley original model and is in line with reports pertaining to impacts of slow Kv1.3 inactivation on adaptive membrane excitability.

**Figure 6:**
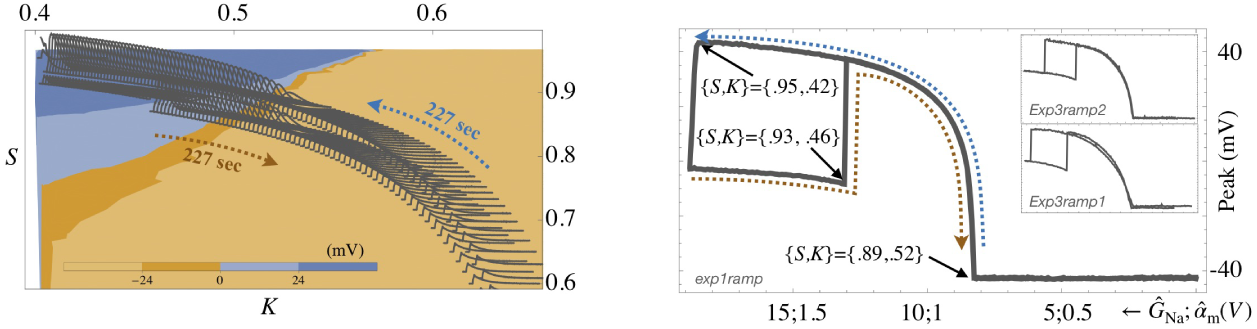
Activity-dependence and hysteretic dynamics due to the impacts of Kv1.3 slow inactivation kinetics. The **left** panel shows membrane responses of a Kv1.3 expressing oocyte as a function of S–K coordinates in a directed walk within the diagram. A gradual (up and down ramp; total trajectory ca. 450 sec) change is implemented in the dynamic clamp parameters of Hodgkin-Huxley sodium conductance (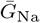 and 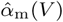). The spikes are plotted over an S–K diagram constructed for that same oocyte. Each trace depicts membrane response to a short above-threshold current stimulation. As the system is moved toward the excitable phase (top-left corner of the diagram), the oocyte responds to the stimulus with a self-propelled depolarization that becomes a fully blown action potential. At some point, as might be expected, the structural exciting force (S) is so high and the kinetic restoring force (K) is so low ({0.95, 0.42}, depicted by arrow in the **right** panel) that the membrane cannot hyperpolarize back to resting potential, and remains stuck in some depolarized not excitable value. Upon return, a clear hysteresis is revealed, reflecting recovery of the Kv1.3 from long lasting inactivation. The insets to the right panel show two repetitions of the same ramp protocol in another oocyte, demonstrating reversibility of the hysteresis phenomenon.

## Concluding remarks

We live in a time marked by capacity to collect data at ever-increasing speed and resolution. As a result, it is tempting to use these data to construct numerical models of increased dimensionality, making them more and more biologically realistic. To avoid the fallacy of attributing importance to each and every measurable parameter, good practice combines methods that point to functional relations between parameters [34] and formulation of low dimensional phase diagrams. But reduction of dimensionality – a *via Regia* to formal understanding – also comes at a price. In many cases it is a unidirectional path where measurables are abstracted and compressed to an extent that loses the explicit properties of the physiological data from the abstract representation. Consequently, once an abstract low dimensional model is constructed, evaluation of impacts and subsequent incorporation of new biological features into the low dimensional model become challenging, if at all possible. Here we approached the problem by implementation of a methodology that has a long and successful history in membrane physiology: system identification using closed-loop control (i.e., voltage-clamp, patch clamp, dynamic clamp). We describe an experimental-theoretical hybrid; a framework enabling bidirectional real-time interaction between abstract low dimensional representation and real biological entities. This is not a post hoc ‘fitting’ procedure; rather, it is a live experiment where the impacts – of a given biological component – on the abstract low dimension representation, are identified by implementing a real-time closed loop design.

Specifically, combining dynamic clamp and heterologous expression of ionic channel proteins in *Xenopus* oocytes, we constructed an excitable system composed of a mix of biologically and computationally expressed components. This experimental configuration enabled systematic sampling of the Hodgkin-Huxley parameter space. The resulting phase diagram validates a theoretically proposed diagram [4], suggesting that maintenance of excitability amidst parametric variation is a low dimensional, physiologically tenable control process. Moreover, we show that the basic ingredients for such control – namely memory and adaptation – are manifested in the phase diagram as a natural outcome of ion channel slow inactivation kinetics.

Many theoretical and experimental analyses show that the wide range of temporal scales involved in slow inactivation is sliced thinly to a degree effectively equivalent to a continuum of scales, indicative of the extensive network of configurations within which the channel protein may “diffuse” giving rise to slow activity-dependent gating and adaptive firing patterns [27, 31, 35–42]. Indeed, slow activity-dependent gating was suggested as a means for maintenance and control of membrane excitability. Specifically, activity dependence of protein kinetics at relatively slow time scales, entailed by multiplicity of protein states, was pointed at as a general “automatic” and local means for stabilization of cellular function, independent of protein synthesis, and operates over a wide – minutes and beyond – range of time scales [3, 4, 43, 44]. Thus, precisely because these ion channels do not have a single, fixed time constant encoded in their molecular structure, but rather slide through multiple states, cells have a built-in mechanism to smoothly function over a larger range of firing patterns and voltages. This partially mitigates the control problem that cells face: getting it right may not require the perfect match between channel numbers that might otherwise be necessary.

Viewed from another angle: multiple states of channel inactivation and recovery from inactivation necessarily result in hysteresis, and the time scales of that hysteresis become a “memory mechanism” [31, 33, 36] so that cells can use it to keep track of their recent pattern of activity and inactivity. This again, expands the time course over which patterns of activity can influence the way the cell responds to physiological inputs. Interestingly, we usually think of the fastest membrane events (action potentials) having little lasting effect on the cells in which they are seen, but looking only at the fast voltage deflections such as action potentials, hides the effects of the slower channel dynamics that influence future events.

It remains to be seen how far the approach described here may be used in system identification of excitable membranes more complicated than the minimal, two conductance single compartment Hodgkin-Huxley configuration. Certainly, cells that contain many different types of ion channels will show a range of time-scales and history-dependence [45]. Developing intuition into how a given set of firing properties depends on conductance densities of many channels may require new kinds of principled dimensionality reduction to complement brute-force numerical simulations.

## Methods

Clusters of *Xenopus* oocytes were kindly provided by N. Dascal’s laboratory (Tel Aviv University). Individual oocytes were separated from their clusters by standard mechanical and enzymatic treatment and kept at 18–20C overnight prior to mRNA injection. The mRNAs were prepared from Kv1.3 carrying vectors kindly provided by Alomone Labs (Jerusalem). Oocytes were allowed to express injected mRNA over 2–6 days before electrophysiological experiments. A two-electrode voltage clamp system (NPI TURBO TEC-03X) and a National Instruments board (NI 625x series) were used to control the experiments. A sufficiently short loop duration (40 *µ*sec) was achieved within Real Time eXpe iment Interface (RTXI; www.rtxi.org) environment implemented in CPP software. The sequence of an experiment was as follows: An oocyte situated in a perfusion chamber was impaled with two Agarose cushioned electrodes prepared as described elsewhere [14]. The bath solution, under continuous perfusion, was composed of 96 mM NaCl, 2 mM KCl, 1 mM MgCl_2_, 1 mM CaCl_2_, 5 mM HEPES, adjusted to pH 7.5 with 5 M NaOH. The reported results are based on experiments conducted at room temperature (air-conditioned ca. 22C) on 7 naive and 13 Kv1.3 injected oocytes. Each experiment began with a voltage-clamp protocol to determine leak conductance, membrane capacitance, potassium Nernst potential and maximal string conductance (in Kv1.3 expressing oocytes). Potassium Nernst potential (−81.5±10.0 mV) was estimated from tail current reversals under voltage clamp. Maximal potassium string conductance (0.23±0.12 mS) was estimated from maximal current and Nernst potential; in several cases, the conductance was also validated using the slope around Nernst potential in tail current protocols (e.g., Figure 3B). Leak conductance in Kv1.3 injected oocytes (0.07±0.06 mS) was estimated from current responses to series of 20 mV hyperpolarizing voltage clamp pulses, from -90 mV holding potential. Cell capacitance in Kv1.3 injected (0.30±0.22 *µ*F) and naive (0.21±0.05 *µ*F) oocytes was estimated from the current step upon instantaneous switch between −2.66 V/sec to +2.66 V/sec (in a range between −70 to −110 mV). Following the voltage-clamp protocol, the system configuration was switched to dynamic clamp mode to screen the S–K plane. A digitally expressed leak conductance was added to set a resting membrane potential around −65 mV; in the case of Kv1.3 injected oocytes, the digital leak added was in most instances two orders of magnitude smaller compared to the biological leak estimated from the above voltage clamp experiments. The actual resting membrane potential upon activation of the dynamic clamp mode was −66 ± 5.4 mV (hereafter, values of the experiment described in Figure 5, in which the dynamic clamp protocol was different, were excluded from the statistics). The average drift in resting potential during a dynamic clamp experiment was 1.5 ± 4.2 mV. For each S;K point, a 99 msec relaxation phase was allowed before a 22 msec duration trace was recorded, within which a 0.5 msec depolarizing stimulus was delivered, followed by 20 msec recording. Above-threshold stimulus amplitude in Kv1.3 injected oocytes varied between cells, ranging from 50 to 60 *µ*A/*µ*F (15.7 ± 13.9 *µ*A), sufficient to reliably elicit a spike in the high-S low-K range. Once set, the stimulus amplitude was kept fixed throughout the experiment. Rate equations of digitally expressed conductances were those used by Hodgkin and Huxley (1952), or – where indicated – the Kv1.3 rate equations [22, 24].

Three comments on difficulties associated with technical aspects of the experimental approach employed here: (1) Completion of a protocol such that S– K coordinates are properly characterized entails voltage clamp procedures for maximal conductance, leak, capacitance, and maintenance of relatively stable resting potential, while going through the S–K plane. Thus, stable electrophysiological settings are necessary for a typical experiment lasting ca. 30 minutes. In our hands, continuous superfusion of the bath medium and use of Agarose cushioned electrodes promoted such stability. (2) Another difficulty arises due to the hardware used. Sometimes, the real time processor and/or the limits of the current injection device were not sufficient to catch-up at high S and low K values, giving rise to “ringing” about the peak of the spike. We assume that advanced hardware can do better. (3) We were not able to implement an experimental condition where sodium conductance was biologically expressed, whereas potassium conductance is computationally expressed. In our hands, the expression of sodium conductance is too weak to support full-blown excitability. Higher expression would have introduced problems in both estimation of maximal conductance and dynamically clamping the fast sodium current, which might be solved by conducting the experiments at lower temperatures.

## Acknowledgement

This work was partially supported by research grants from the Leir Foundation (EM, SM) the National Institutes of Health (EM), and the Israel Science Foundation (SM). The authors thank Leonid Oddesky, Tamar Galateano and Yael Abuchatzera for technical support, Nathan Dascal’s group (Tel Aviv University) for generous supply of oocytes, Alomone Labs (Jerusalem) for channel carrying vectors, and Omri Barak, Erez Braun, Daniel Dagan and Michele Giugliano for helpful comments.

